# Antagonist muscle activity during reactive balance responses is elevated in Parkinson’s disease and in balance impairment

**DOI:** 10.1101/343004

**Authors:** Kimberly C. Lang, Madeleine E. Hackney, Lena H. Ting, J. Lucas McKay

**Affiliations:** Graduate Division of Biological and Biomedical Sciences, Emory University, Atlanta, Georgia; Division of General Medicine and Geriatrics, Emory University School of Medicine, Atlanta, Georgia; Rehabilitation R&D Center, Atlanta VA Medical Center, Atlanta, Georgia; Department of Rehabilitation Medicine, Division of Physical Therapy, Emory University, Atlanta, Georgia; The Wallace H. Coulter Department of Biomedical Engineering at Emory University and Georgia Tech, Atlanta, Georgia

**Keywords:** Parkinson’s disease, co-contraction, modulation, reactive balance, antagonist activation

## Abstract

BACKGROUND: Abnormal antagonist leg muscle activity could indicate increased muscle co-contraction and clarify mechanisms of balance impairments in Parkinson’s disease (PD). Prior studies in carefully selected patients showed PD patients demonstrate earlier, longer, and larger antagonist muscle activation during reactive balance responses to perturbations. RESEARCH QUESTION: Here, we tested whether antagonist leg muscle activity was abnormal in a group of PD patients who were not selected for phenotype, and most of whom had volunteered for exercise-based rehabilitation. METHODS: We compared antagonist activation during reactive balance responses to multidirectional support-surface translation perturbations in 31 patients with mild-moderate PD (age 68±9; H&Y 1-3; UPDRS -III 32±10) and 13 matched individuals (age 65±9). We quantified modulation of muscle activity (i.e., the ability to activate and inhibit muscles appropriately according to the perturbation direction) using modulation indices (MI) derived from minimum and maximum EMG activation levels observed across perturbation directions. RESULTS: Antagonist leg muscle activity was abnormal in unselected PD patients compared to controls. Linear mixed models identified significant associations between impaired modulation and PD (P<0.05), PD severity (P<0.01), and balance ability (P<0.05), but not age (P=0.10). SIGNIFICANCE: Antagonist activity is increased during reactive balance responses in PD patients of varying phenotypes who are candidates for rehabilitation. Abnormal antagonist activity may contribute to balance impairments in PD and be a potential rehabilitation target or outcome measure.

**Funding Sources:** This work was supported by the National Institutes of Health (NIH) KL2 TR000455, K25 HD086276, R01 HD46922, R21 HD075612, TL1TR000456, and UL1 TR000424. The study sponsors had no role in study design; in the collection, analysis and interpretation of data; in the writing of the report; or in the decision to submit the article for publication.

**Declaration of Interest:** Conflicts of interest: None.

**Author Contributions:** Research project: Conception, JLM, LHT, MEH; Organization, JLM, KCL, LHT, MEH; Execution, JLM, KCL, LHT, MEH.

Statistical Analysis: Design and Execution, JLM, KCL; Review and Critique, LHT, MEH. Manuscript Preparation: Writing of the first draft, KCL; Review and Critique: JLM, LHT, MEH.

**Highlights:**

- We quantified abnormal antagonist muscle activation during balance in PD.
- Abnormalities were related to PD severity as well as to overall balance ability.
- Abnormalities were similar across PD phenotypes with and without tremor.
- Balance antagonist activation may be a useful rehabilitative outcome measure.

## 1 Introduction

Abnormal antagonist muscle activity can cause joint stiffening by concurrently activating paired agonist and antagonist muscles (“co-contraction” or “co-activation”) [1-4], which may contribute to balance impairment in people with Parkinson’s disease (PD). Prior studies in PD patients carefully selected for postural difficulties and minimal tremor [5, 6] demonstrate earlier, longer, and larger antagonist muscle activation during reactive balance responses to support surface perturbations compared to controls [5-8]. Evaluation of antagonist muscle activation during balance could therefore potentially inform improved rehabilitative outcome measures (e.g., [9]). However, it is unclear whether antagonist muscle activity during reactive balance responses is abnormal in PD patients who are not selected by phenotype and are candidates for exercise-based rehabilitation.

While co-contraction is not a PD-specific phenomenon [10], its elevation with age and PD and its effects on functional balance make it relevant to understanding balance impairment in PD. In adults without PD, muscle co-contraction is associated with functional changes in behavior, including increased sway [11-14], increased risk of falls [15, 16], and decreased functional reach distance and functional stability boundaries [13].

Here, our objective was to determine whether antagonist muscle activity during balance responses was increased across leg muscles in people with PD who were not selected by phenotype. We recorded automatic postural responses induced by multidirectional translational support surface perturbations and examined subsequent muscle activation [17, 18] in PD patients and matched neurotypical individuals. As an assay of abnormal antagonist activity, we quantified the ability to activate and inhibit muscles appropriately according to the perturbation direction using modulation indices (MI) derived from minimum and maximum EMG levels observed across directions. Our primary analyses examined associations between the presence of PD and decreased modulation. To clarify the role of other predictors, we also performed secondary analyses to assess the associations between decreased muscle modulation and 1) age, 2) interaction between PD and age, 3) balance ability, 4) PD phenotype, and 5) PD severity.

## 2 Methods

### 2.1. Participants

We performed a cross-sectional observational study using baseline measures from a longitudinal study of exercise-based rehabilitation. PD patients (n=34) and age-matched individuals without PD (“nonPD,” n=16) were recruited from the Atlanta area from December 2013 through May 2017. Among PD patients, the majority (21/34) were enrolled into a two-arm randomized trial with dance-based exercise rehabilitation and non-contact control arms; the remaining patients and all matched individuals were allocated directly to the non-contact control arm. No screening on symptom phenotype was performed. Participants provided written consent according to protocols approved by the Institutional Review Boards of Emory University and/or the Georgia Institute of Technology.

Inclusion criteria were: age ≥ 35, vision corrected if necessary, ability to walk ≥ 10 feet with or without an assistive device, normal perception of vibration and light touch on feet, no dance class participation within the previous 6 months, and demonstrated response to levodopa (PD only). Exclusion criteria were: significant musculoskeletal, cognitive, or neurological impairments other than PD as determined by the investigators.

After enrollment, participants were excluded from analysis for the following reasons: neurological diagnosis other than PD disclosed after study entry (N=1 PD, N=1 nonPD), noncompliance with OFF medication state (N=1 PD), inability to complete reactive balance protocol (N=1 PD), suspected undiagnosed cognitive impairment (N=1 nonPD), and technical difficulties in data processing (N=1 nonPD).

### 2.2. Assessment protocol

All participants were assessed according to a standardized protocol that spanned 3-4 hours including informed consent, collection of clinical and demographic information, and assessment of clinical and reactive balance. PD symptom severity was assessed by the Unified Parkinson’s Disease Rating Scale III (UPDRS-III) [19], by a Movement Disorders Society-certified rater (MEH) either in-person or on video. PD phenotype (tremor dominant, TD; indeterminate, ID; postural instability and gait difficulty, PIGD) was calculated using standard formulae [20]. Balance ability was assessed with the Fullerton Advanced Balance Scale (FAB) [21] and Berg Balance Scale (BBS) [22]. Gait was assessed with the Dynamic Gait Index (DGI) [23]. Freezing of gait was assessed with the Freezing of Gait Questionnaire B (FOGQ-B) [24]. All PD patients were assessed in the 12-hour OFF medication state.

### 2.3. Reactive balance assessments

Participants stood on a custom perturbation platform that produced ramp-and-hold support-surface translations (7.5 cm peak displacement, 15 cm/s peak velocity, 0.1 g peak acceleration) [9]. Feet were positioned parallel with medial aspects 28 cm apart and arms were crossed across the chest. Participants experienced 3 forward perturbations of the support surface to reduce startle (or “first-trial”) effects before being tested with a set of 36 randomized perturbations in 12 evenly distributed horizontal-plane directions (Figure 1). Perturbation trials that elicited stepping responses were repeated at the end of the randomized block if possible.

**Figure 1.**
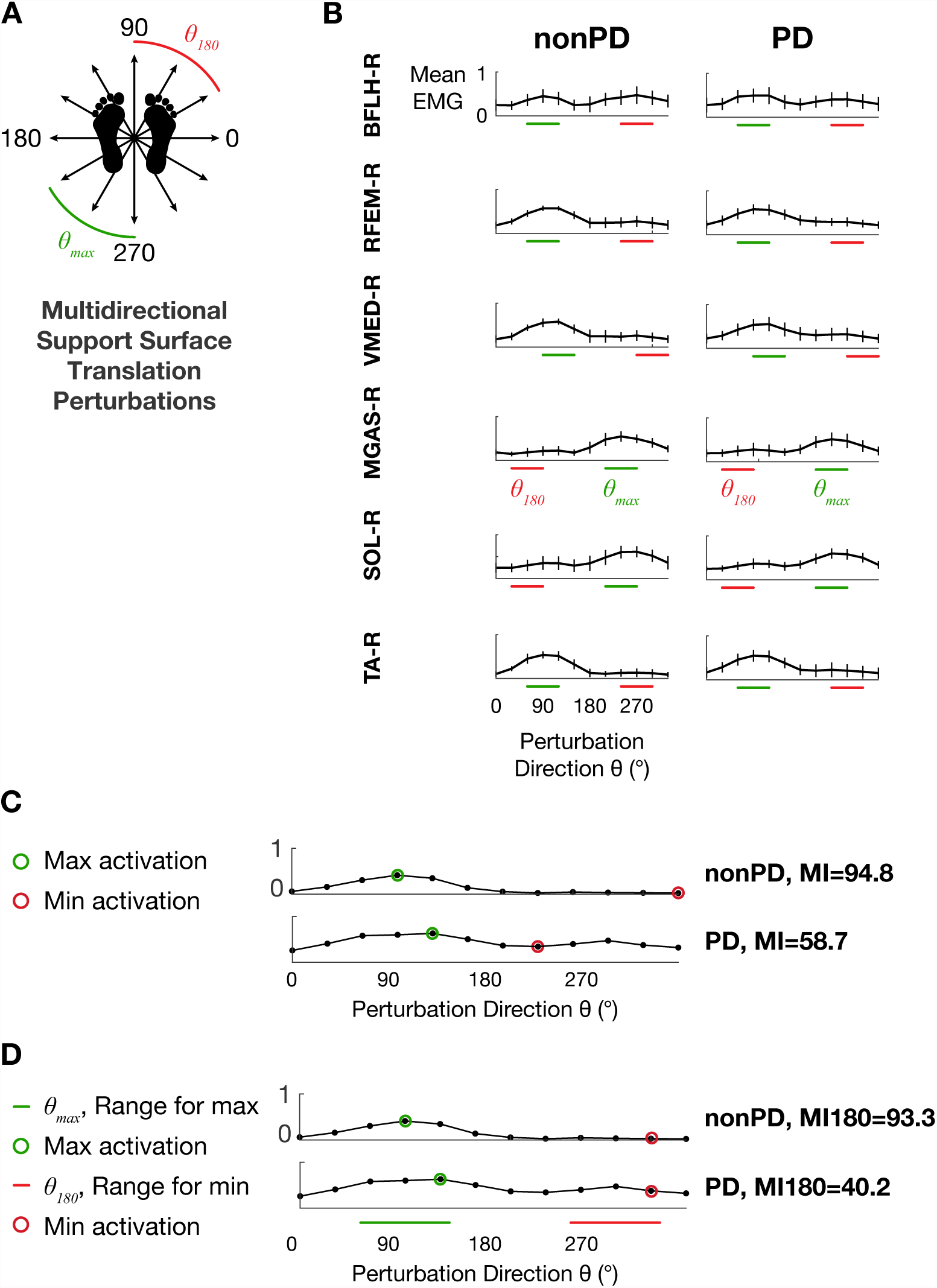
A: Schematic depiction of multidirectional support surface translation perturbations. Green and red perturbation directions correspond to those for which maximum values were observed most frequently for MGAS-R and those directly opposite (see D). B: Tuning curves from the nonPD and PD groups depicting mean EMG activity during the APRX time bin (70-450 ms after perturbation onset). Horizontal bars indicate perturbation direction ranges *θmax* and *θ_180_* used for calculation of modulation index MI180. C. Examples of calculation of MI (Equation 1) and MI180 (Equation 2) for TA from two different participants.

### 2.4. EMG processing

Surface EMG activity was collected from 11 lower limb muscles: bilateral *soleus* (left, SOLL; right, SOL-R), *medial gastrocnemius* (MGAS-L, MGAS-R), *tibialis anterior* (TA-L, TA-R), *biceps femoris long head* (BFLH-L, BFLH-R), *rectus femoris* (RFEM-L, RFEM-R) and right *vastus medialis* (VMED-R). Silver/silver chloride disc electrodes were placed 2 cm apart at the motor point [25]. EMG data were recorded using telemetered EMG (Konigsburg, Pasadena, CA) and synchronized to kinematic data (120 Hz) using Vicon motion capture equipment (Oxford Metrics, Denver, CO). EMG data were recorded at either 1080 or 1200 Hz depending on the equipment version. EMG recordings were processed offline (high-pass, 35 Hz, de-mean, rectify, low-pass, 40 Hz) [9]. Trials eliciting stepping responses or spotter intervention were identified in video records and excluded from analyses. Trials with significant EMG motion artifacts were identified by visual inspection and excluded from analyses. After exclusions due to steps or EMG quality concerns, the number of trials available per perturbation direction per participant ranged from 0 to 5 with an average of 3.0 ± 0.3.

### 2.5. Muscle activity modulation indices: MI and MI180

In order to assess modulation of muscle activity during reactive balance, we computed a muscle “modulation index” that described the ability to activate and inhibit each muscle appropriately according to the perturbation direction. Because of the increased number of experimental conditions compared to previous studies, we developed two extensions of an existing modulation index that was initially developed to assess antagonist activity in only two movement directions [18]. In previous work, the movement directions that require each muscle to be activated (as an agonist) or inhibited (as an antagonist) were obvious from the biomechanical constraints of the task. In the multidirectional perturbation protocol used here, each muscle exhibits a continuum of activity from agonist to antagonist as a function of perturbation direction.

Therefore, we calculated mean EMG levels during two time bins within each trial that encompassed the medium- and medium- and long-latency automatic postural response: 100- 175 ms (APR1) and 70-450 ms (APRX) after perturbation onset [5], and subsequently assembled mean APR1 and APRX EMG levels into tuning curves that described muscle activity as a function of perturbation direction (Figure 1). Then, we used the maximum and minimum values of each tuning curve for each muscle for each participant to compute the modulation index (MI) using the following equation (Figure 1):
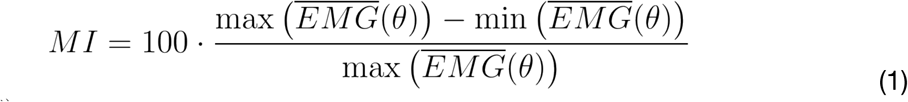

where 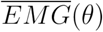 indicates the vector of 12 mean EMG values for the 12 perturbation directions.

While the MI value reflects the greatest amount of modulation across the 12 perturbation directions, in some cases, it did not capture abnormally elevated activity 180° from the perturbation direction for which the muscles were maximally activated, and in which the muscles could reasonably be assumed to be antagonists due to the biomechanical constraints of the task. Therefore, we developed a similar formula to calculate a more physiologically-relevant index (MI180), in which the maximum value of each tuning curve was identified within the range *θ_max_* of the 3 perturbation directions for which maximum EMG values were observed most frequently (Figure 1) and the minimum value was identified within the range *θ_180_* directly opposite *θ_max_*:
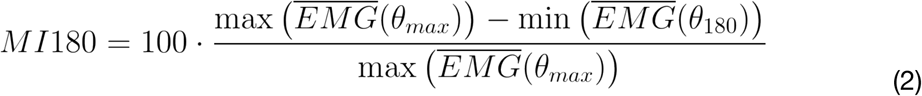

where 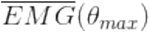 indicates the vector of 3 mean EMG values for the 3 perturbation directions included in *θ_max_*, and 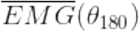 corresponds similarly to the vector of 3 mean EMG values for *θ_180_*.

### 2.6. Statistical Analysis

Differences between the PD and nonPD groups in demographic and clinical variables were assessed with chi-square tests and independent samples *t*-tests as appropriate.

For each muscle recorded, separate chi-square tests of homogeneity were performed to assess crude differences in modulation between participants with vs. without PD, between participants above vs. below the sample median in age, and between participants above vs. below the sample median in balance ability, as assessed by FAB. For these tests, modulation indices (MI and MI180 in both APR1 and APRX) were dichotomized about median values.

Associations between predictors (PD, age above the sample median, and balance ability below the sample median) and the presence of MI below the median were expressed as odds ratios (OR) ± 95% CI. OR>1 indicate strong associations between the presence of a given predictor and the presence of low modulation. Primary analyses were conducted with MI in APRX (detailed below) and repeated with MI in APR1 and MI180 in APR1 and APRX.

To estimate the association between study variables and modulation across muscles, multivariate linear regression analyses were used to examine the effects of predictors of interest, including PD, age, balance, PD severity, PD phenotype, and the interaction between PD and age.

To test whether the presence of PD was associated with muscle modulation, we fit the following linear mixed model:
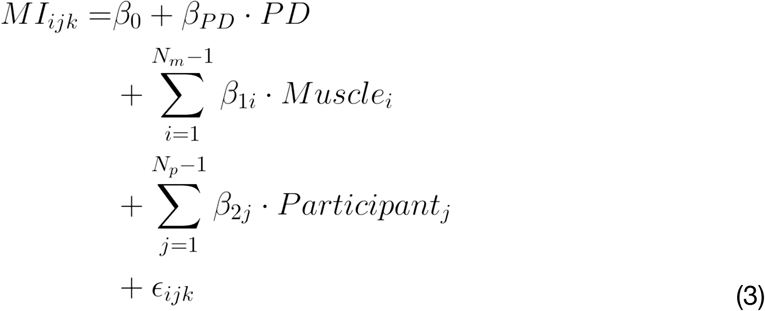

in order to evaluate the following null hypothesis with an *F* test:

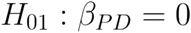

In Equation 3, the indicator variable *PD* is 1 for participants with PD and 0 otherwise, *β_1i_* is the beta coefficient for the fixed effect of muscle *i* (with TA as the reference group) and *β_2j_* is the beta coefficient for the random effect of participant *j*.

To test whether age was associated with muscle modulation, we fit the following model:
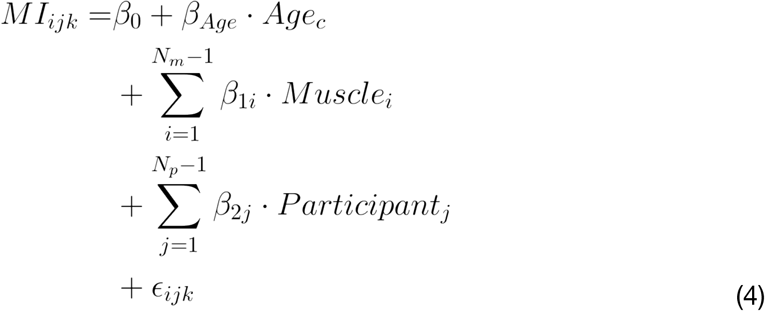

where *Age_c_* designates participant age centered about the sample median, and evaluated the null hypothesis:

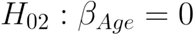

Similarly, to test whether balance ability as measured by FAB was associated with muscle modulation, we fit the following linear mixed model:
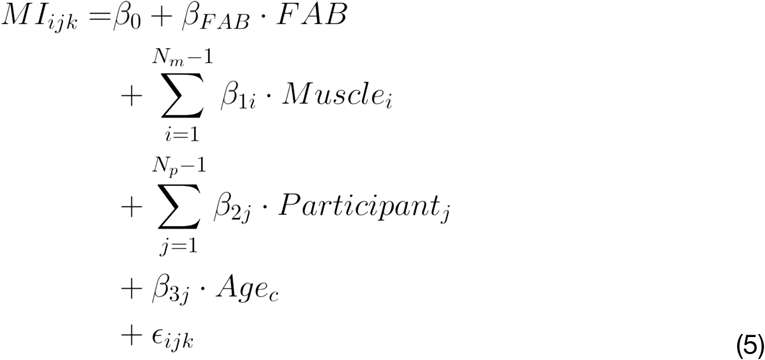

where *FAB* designates total FAB score, and the following null hypothesis was evaluated with an *F* test:

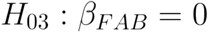

Additional linear mixed models evaluating associations between additional candidate predictor variables and modulation are presented in Supplementary Information 1.

## 3 Results

### 3.1. Participant characteristics

Demographic and clinical characteristics of the study participants are presented in Table 1. No significant differences were observed between the PD and nonPD groups in sex, age, or BMI. Compared to nonPD, the PD group had significantly poorer balance performance on FAB, BBS, and DGI (all P values<0.01), and significantly increased prevalence of previous falls (P=0.03).

**Table 1.** Demographic and clinical characteristics of study participants with and without Parkinson’s disease (PD).

|  | PD<br>(N=31) | nonPD<br>(N=13) | P Value |
| --- | --- | --- | --- |
| Sex (N, %) |  |  | 0.60 |
| Male | 17, 55% | 6, 46% |  |
| Female | 14, 45% | 7, 54% |  |
| Age, y, mean±SD | 67.6 ± 8.8 | 64.5 ± 8.8 | 0.28 |
| BMI, kg/m<sup>2</sup>, mean±SD | 25.6 ± 4.0 | 26.0 ± 3.8 | 0.76 |
| Behavioral balance measures |  |  |  |
| BBS (0-56), mean±SD | 52.2 ± 4.4 | 55.1 ± 1.3 | <0.01* |
| FAB (0-40), mean±SD | 29.2 ± 5.7 | 33.1 ± 3.1 | <0.01* |
| DGI (0-24), mean±SD<sup>a</sup> | 20.0 ± 3.5 | 22.5 ± 1.3 | <0.01* |
| Fall History |  |  | 0.03* |
| 0 falls in previous 12 months, (N, %) | 13, 42% | 10, 77% |  |
| ≥1 fall in previous 12 months, (N, %) | 18, 58% | 3, 23% | |
| PD clinical features |  |  |  |
| PD duration, y, mean±SD | 7.5 ± 5.9 | - | |
| UPDRS-III (0-108), mean±SD | 31.7 ± 9.5 | - | |
| UPDRS items, mean±SD | | - | |
| Leg rigidity (III.22, 0-8) | 1.9 ± 2.0 | - | |
| Posture (III.28, 0-4) | 1.0 ± 1.0 | - | |
| Gait (III.29, 0-4) | 1.1 ± 0.6 | - | |
| Postural stability (III.30, 0-4) | 0.8 ± 0.7 | - | |
| Modified Hoehn & Yahr Stage, (N, %) |  | - |  |
| 1 | 1, 3% | - |  |
| 1.5 | 5, 16% | - |  |
| 2 | 13, 42% | - |  |
| 2.5 | 4, 13% | - |  |
| 3 | 8, 26% | - |  |
| PD phenotype, (N, %) |  | - |  |
| Postural Instability and Gait Disability (PIGD) | 19, 61% | - |  |
| Indeterminate (ID) | 3, 10% | - |  |
| Tremor-Dominant (TD) | 9, 29% | - |  |
| Freezing of Gait, (N, %)<sup>b</sup> |  | - |  |
| Freezer | 14, 45% | - |  |
| Non-freezer | 15, 48% | - |  |

Abbreviations: BBS, Berg Balance Scale; FAB, Fullerton Advanced Balance Scale; DGI, Dynamic Gait Index. <sup>a</sup>PD N=29. <sup>b</sup>PD N=29. \*P<0.05.

### 3.2. Description of muscle activity across perturbation directions

Tuning curves exhibited clear cosine tuning (Figure 1) consistent with those reported previously in the literature [5, 26]. Average APRX tuning curve widths at half maximum [27] were 115±7° and 111±11° for the nonPD and PD groups, respectively. Across all subjects and muscles, mean values of modulation indices in APRX were 71.9±12.9, 36.4-96.6 (mean±SD, range) for MI and 59.4±31.8, -301.6-94.8 for MI180. In APR1, the values were 70.8±15.2, 20.7- 98.4 (MI) and 63.1±24.2, -160.3-98.4 (MI180). Negative values observed in MI180 corresponded to tuning curves in which muscles were more strongly activated in the *θ_180_* range of perturbation directions and accounted for a small percentage of tuning curves in both the PD (2.4% in APRX, 1.2% in APR1) and nonPD groups (3.5% in APRX, 0.7% in APR1).

### 3.3. PD, age, and impaired balance ability were associated with impaired modulation in some individual muscles

Univariate analyses showed that PD was associated with lower MI for each muscle analyzed during the APRX time window (Figure 2A, filled circles; note that all Odds Ratios [OR] >1). This association was statistically significant for TA (OR=4.02, P<0.01). PD was associated with lower MI180 in 4/6 muscles analyzed during APRX (Figure 2A, unfilled circles). Age was also associated with lower MI in APRX (OR: 2.79±1.67, range 1.21-5.69), particularly for BFLH (P<0.01), SOL (P<0.05), and TA (P<0.001) (Figure 2B). Low FAB score was associated with lower MI for both BFLH (OR: 2.52, 95% CL: 1.07-5.95, P=0.03) and TA (OR: 4.59, 95% CL: 1.87-11.26, P<0.001) during APRX. Analyses during APR1 showed inconsistent associations between PD and impaired modulation (significant in SOL with MI180 (OR: 3.12 [1.18-8.25], P=0.02); Figure 2A).

**Figure 2.**
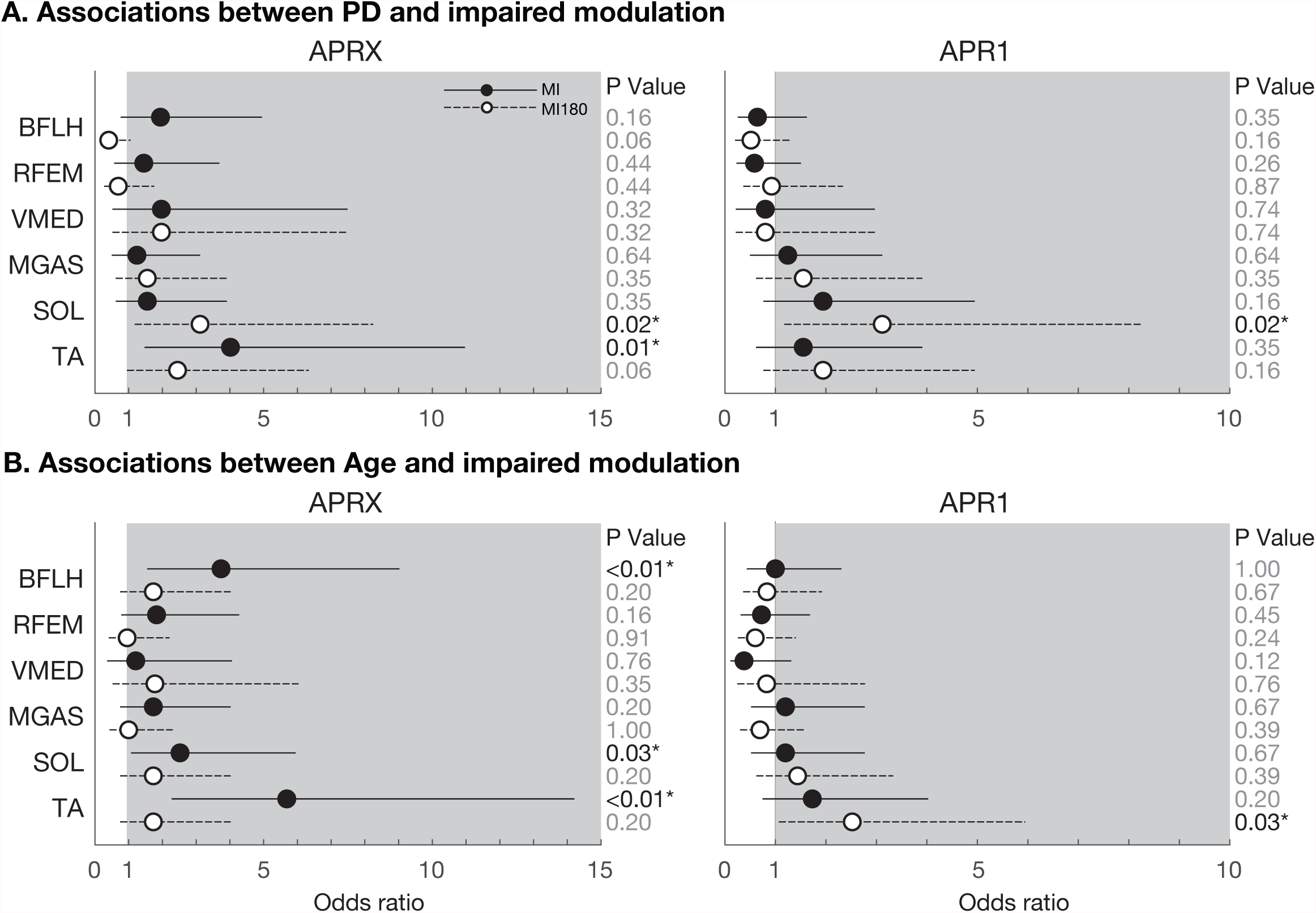
Associations between PD (A) and Age (B) and impaired modulation in analyses of individual muscles. Associations are described as Odds Ratios (OR) calculated separately using both MI and MI180 modulation indices derived from both APR1 and APRX time bins. Solid lines and dots represent the OR and 95% confidence limits for modulation index MI; dashed lines and open dots represent modulation index MI180. Odds ratios > 1 (shaded area) indicate that the presence of the risk factor (PD or Age) is strongly associated with the presence of impaired modulation for that muscle.

### 3.4. PD, PD severity, and impaired balance ability were associated with impaired modulation across muscles

Across muscles, linear mixed models identified significant associations between PD (P<0.05) and PD severity (P<0.01) and decreased MI during APRX (Table 2). Higher FAB score was significantly associated with increased MI during APRX (P<0.05). There was only marginal evidence of an association between increased age and decreased MI (P=0.10), or, similarly, for interaction between PD and age in the effect on MI (P=0.13). Linear mixed models that stratified the PD group by PD phenotype identified strong associations between each phenotype (TD, ID, and PIGD) and decreased MI although identified parameters were only marginally significant (P=0.06, 0.05, 0.15). Associations between these predictor variables and MI180 were the same in direction but decreased in magnitude by ≈34%. The only exception to this pattern was that no association was identified between FAB score and MI180. No significant associations between predictors and modulation indices were identified in analyses of APR1 (Table S1).

**Table 2.** Associations between predictors of interest and muscle modulation indices MI and MI180.

| Predictor | MI |  |  | MI180 |  |  |
| --- | --- | --- | --- | --- | --- | --- |
| | $\beta$ | 95% CI | P Value | $\beta$ | 95% CI | P Value |
| PD | -4.26 | -8.31, -0.21 | 0.04* | -3.34 | -11.66, 4.98 | 0.43 |
| Age | -0.18 | -0.40, 0.03 | 0.10 | -0.14 | -0.58, 0.30 | 0.53 |
| FAB | 0.38 | 0.005, 0.75 | <0.05* | -0.04 | -0.83, 0.74 | 0.91 |
| PD Severity | -0.16 | -0.26, -0.05 | <0.01* | -0.06 | -0.29, 0.18 | 0.64 |
| PD Phenotype |  |  |  |  |  |  |
| PIGD | -3.25 | -7.67, 1.18 | 0.15 | -2.41 | -11.64, 6.82 | 0.61 |
| TD | -5.22 | -10.55, 0.12 | 0.06 | -3.85 | -15.00, 7.29 | 0.50 |
| ID | -7.82 | -15.69, 0.04 | 0.05 | -7.70 | -24.11, 8.71 | 0.36 |
| PD•Age | -0.36 | -0.83, 0.10 | 0.13 | -0.09 | -1.09, 0.91 | 0.86 |

\*p<0.05. Abbreviations: FAB, Fullerton Advanced Balance Scale; PIGD, Postural Instability and Gait Difficulty; TD, Tremor-Dominant; ID, Indeterminate. Mixed model results reflect the APRX time window.

## 4 Discussion

This study’s main result was that leg muscle activity during reactive balance was abnormal in a group of mild-moderate PD patients with a range of symptom phenotypes. We found that lower muscle modulation across perturbation directions – an estimate of an impaired ability to appropriately inhibit muscles according to the biomechanical requirements of the balance task – was predicted by the presence of PD and by PD severity, and that, importantly, these findings were common across the TD, PIGD, and indeterminate phenotypes. Overall, these results extend previous seminal studies in carefully selected PD patients and provide additional evidence that antagonist muscle activation could be a useful rehabilitative target.

Our main motivation for this study was to test whether the results of key earlier studies held among patients who were representative of those interested in rehabilitation and not selected on phenotype [5, 6]. Prior work identified abnormalities during reactive balance muscle activity in PD patients selected for “gait and postural abnormalities” [6] or “axial and/or postural problems and minimal tremor” [5]. While carefully selecting patients decreases variability and is clearly appropriate for foundational research, we propose that it is critical to establish that the results generalize to rehabilitation, where restricting enrollment to certain patient subgroups is typically impractical and uncommon.

Importantly, while we anticipated differences between the PIGD and TD phenotypes on the balance task (e.g., potentially no association between TD parkinsonism and abnormal balance muscle activity), we found that compared to the overall PD effect on MI (*β*=-4.26), the effects of each particular PD phenotype on MI were relatively similar, ranging from only moderate attenuation (PIGD, *β*=-3.25, attenuation of overall PD effect of -24%) to substantial strengthening (indeterminate, *β*=-7.82, +84%) of the overall PD effect.

While comparing our nonPD group to those of previous studies is difficult – there are no obvious clinical variables to use – it is encouraging that the prevalence of previous falls in our nonPD group recruited from the metro Atlanta area (23%) was similar to that reported among the spouses of PD patients in the Netherlands (27% [28]). This provides some evidence that the neurotypical nonPD group here is comparable to those recruited from other geographic regions (i.e., Washington and Oregon [5, 6, 8], Western Europe [7]) with different sociodemographic profiles.

One important limitation to note is that although we examined a larger sample of patients (n=31) than many studies (n=9-13 patients [5-7, 9]), sample size limitations prevented us from imposing the most stringent phenotype classification criteria that are currently recommended [29]. It remains to be seen whether the associations between phenotypes and modulation reported here would be affected by the use of more stringent criteria. However, based on the strong associations with impaired modulation observed in all phenotype groups, we believe it to be unlikely.

We were surprised that age was not significantly associated with muscle modulation here, given that co-contraction is elevated in neurotypical older adults compared to young adults [10]. We speculate that including college-aged participants would probably have resulted in a clear age effect, although potentially one that was nonlinear with time, given that we did not observe a strong effect of age in this sample, which ranged from 39-86.

The presence of a significant association between FAB score and muscle modulation supports the idea that outcome measures derived from antagonist muscle activation could be useful in the general geriatric population, although more studies are required to confirm this. Reports suggest that training may reduce co-contraction during postural control in neurotypical older adults [30] and PD [9]. However, in the linear mixed model used here (Equation 5) sample size prevented us from controlling for the presence of PD. The identified association may in part reflect a PD effect rather than a balance effect per se.

From a methodological perspective, the modulation indices developed here may be useful in contexts other than reactive balance for capturing muscle modulation without requiring pre-specified directions of agonist and antagonist activity. These results offer a measure of the greatest possible amount of modulation (MI) and a measure of the amount of modulation that occurs when effective agonist activity is important for reactive balance (MI180). MI180 also captures instances of antagonist activity that are greater than agonist activity, which were infrequent here.

In summary, we found evidence that the presence of PD, PD severity, and reduced balance ability were related to a measure of elevated leg muscle antagonist activity during reactive balance. It remains to be seen whether abnormal muscle activity results from primary PD disease processes or represents a compensatory strategy (adaptive or maladaptive). However, our findings suggest that there is a relationship between antagonist activity and balance impairment in PD that generally holds for the TD and PIGD phenotypes. Consequently, elevated antagonist activity and the resulting co-contraction could be a useful target or outcome measure for balance rehabilitation.

## Supplementary Information 1

## 1. Additional linear mixed models

In addition to the linear mixed models described in the main text, we fit the following linear mixed models in order to evaluate associations between additional candidate predictor variables and muscle modulation.

### 1.1. Interaction between PD and age

To test whether associations between PD and modulation were modified by age, we fit the following linear mixed model with an interaction term:S
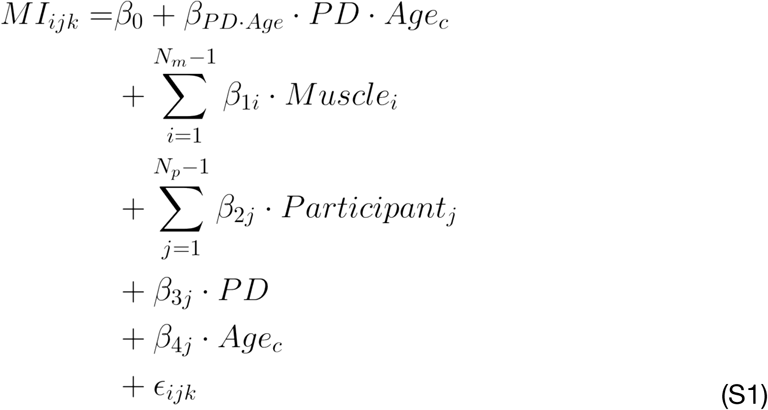

with the following null hypothesis:

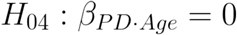

### 1.2. PD phenotype

To test whether phenotype (TD, ID, PIGD, nonPD) was associated with MI modulation during APRX across all muscles, we fit the following linear mixed model, with variables as defined in the main text:S
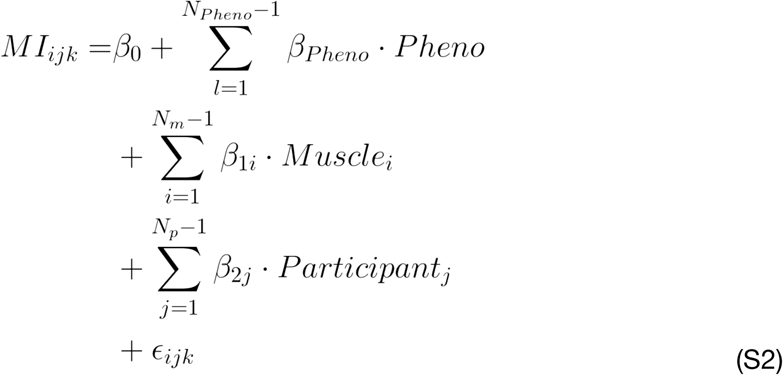

where *β_Pheno_* refers to the beta coefficient for phenotype l, with nonPD as the reference group. The following null hypothesis was evaluated with a Type III F-test:

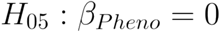

### 1.3. PD severity

To test whether PD severity (UPDRS-III score) was associated with MI modulation during APRX across all muscles, we fit the following linear mixed model:S
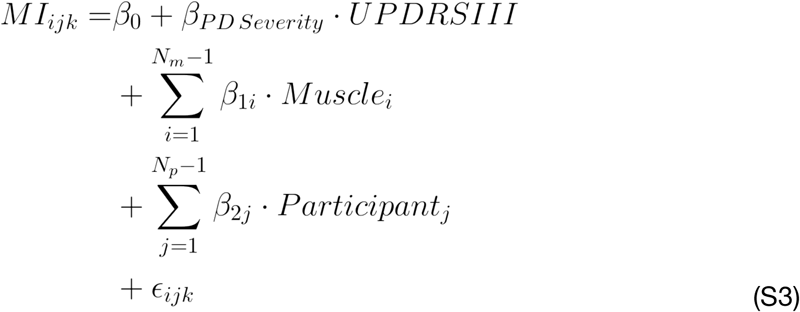

where *β_PDSeverity_* refers to the beta coefficient for UPDRS-III score. The following null hypothesis was evaluated with a Type III F-test:

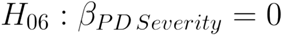

## 2. Associations between study variables and modulation indices in APR1

Across muscles, linear mixed models identified no significant associations between predictors and either modulation index in the APR1 time bin (Table S1).

**Table S1.**
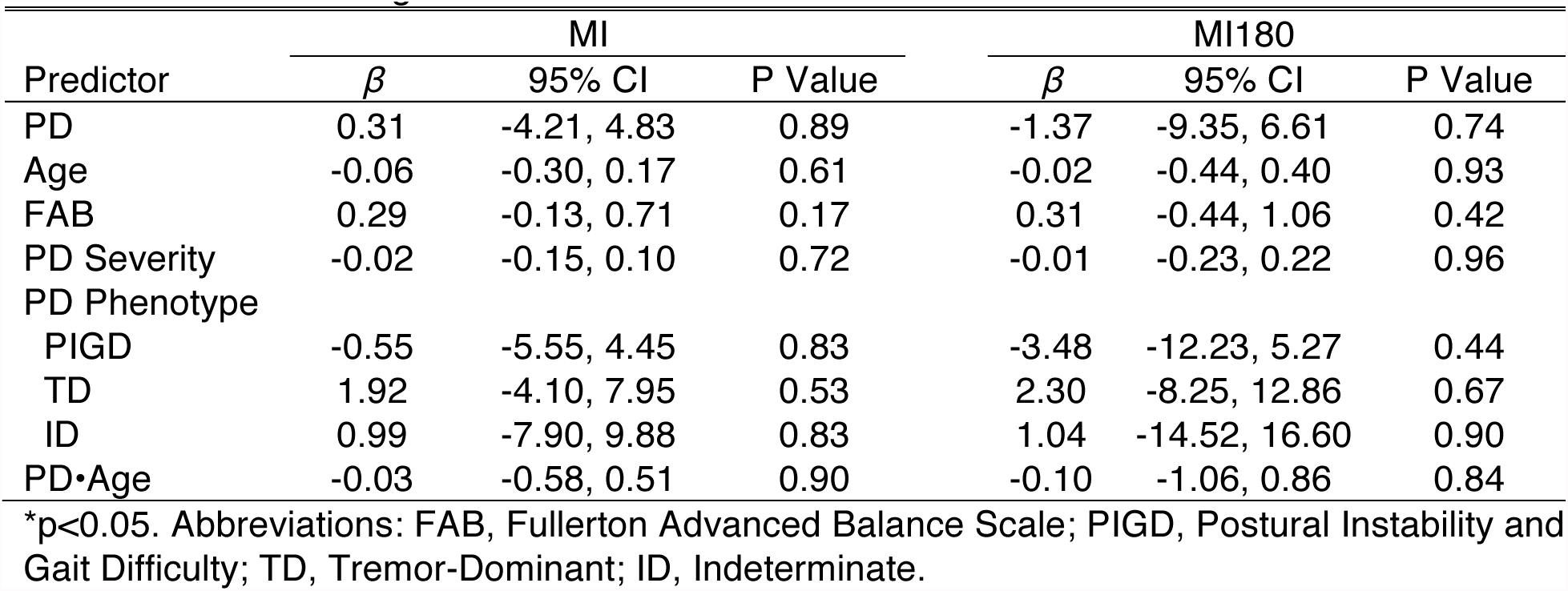
Associations between predictors of interest and muscle modulation indices MI and MI180 calculated during the APR1 time window.

| Predictor | MI |  |  | MI180 |  |  |
| --- | --- | --- | --- | --- | --- | --- |
| | $\beta$ | 95% CI | P Value | $\beta$ | 95% CI | P Value |
| PD | 0.31 | -4.21, 4.83 | 0.89 | -1.37 | -9.35, 6.61 | 0.74 |
| Age | -0.06 | -0.30, 0.17 | 0.61 | -0.02 | -0.44, 0.40 | 0.93 |
| FAB | 0.29 | -0.13, 0.71 | 0.17 | 0.31 | -0.44, 1.06 | 0.42 |
| PD Severity | -0.02 | -0.15, 0.10 | 0.72 | -0.01 | -0.23, 0.22 | 0.96 |
| PD Phenotype |  |  |  |  |  |  |
| PIGD | -0.55 | -5.55, 4.45 | 0.83 | -3.48 | -12.23, 5.27 | 0.44 |
| TD | 1.92 | -4.10, 7.95 | 0.53 | 2.30 | -8.25, 12.86 | 0.67 |
| ID | 0.99 | -7.90, 9.88 | 0.83 | 1.04 | -14.52, 16.60 | 0.90 |
| PD•Age | -0.03 | -0.58, 0.51 | 0.90 | -0.10 | -1.06, 0.86 | 0.84 |

\*p<0.05. Abbreviations: FAB, Fullerton Advanced Balance Scale; PIGD, Postural Instability and Gait Difficulty; TD, Tremor-Dominant; ID, Indeterminate.

